# Spenito-dependent metabolic sexual dimorphism intrinsic to fat storage cells

**DOI:** 10.1101/2023.02.17.528952

**Authors:** Arely V. Diaz, Tyler Matheny, Daniel Stephenson, Travis Nemkov, Angelo D’Alessandro, Tânia Reis

## Abstract

Metabolism in males and females is distinct. Differences are usually linked to sexual reproduction, with circulating signals (e.g. hormones) playing major roles. By contrast, sex differences prior to sexual maturity and intrinsic to individual metabolic tissues are less understood. We analyzed *Drosophila melanogaster* larvae and find that males store more fat than females, the opposite of the sexual dimorphism in adults. We show that metabolic differences are intrinsic to the major fat storage tissue, including many differences in the expression of metabolic genes. Our previous work identified fat storage roles for Spenito (Nito), a conserved RNA-binding protein and regulator of sex determination. Nito knockdown specifically in the fat storage tissue abolished fat differences between males and females. We further show that Nito is required for sex-specific expression of the master regulator of sex determination, Sex-lethal (Sxl). “Feminization” of fat storage cells via tissue-specific overexpression of a Sxl target gene made larvae lean, reduced the fat differences between males and females, and induced female-like metabolic gene expression. Altogether, this study supports a model in which Nito autonomously controls sexual dimorphisms and differential expression of metabolic genes in fat cells in part through its regulation of the sex determination pathway.

## Introduction

Males and females differ fundamentally with regard to metabolism [1], but the underlying molecular mechanisms regulating these differences are incompletely understood. Most studies focus on the importance of sex chromosomes and sex hormones on regulating these differences, especially how signals from the gonads influence metabolism in other tissues (e.g. estrogen) [2]. Less is known about the effects of sex chromosome constitution in tissues not directly involved in sexual reproduction, and to what extent these differences contribute to the sexual dimorphisms observed at the organismal level, including metabolic dimorphism [3].

Metabolic dimorphism is well documented in sexually mature adult *Drosophila melanogaster* (reviewed in [4]). Differences in triglyceride storage and breakdown [5-7], lipid composition [8], and obesogenic responses to diets [9,10] have all been identified. Furthermore, dietary switches can affect males and females differently [11-13]. Some behaviors that change metabolic outcomes are also dimorphic: feeding and locomotor activities differ between the sexes in certain dietary and physiological conditions across the lifespan [14-17]. As in humans, most metabolic dimorphisms in flies have been linked to circulating signals between different tissues. Sex peptide is a hormone found in sperm that influences female physiology and behavior after mating, including feeding [18] and nutrient utilization [19]. Sex differences in adult *Drosophila* courtship behaviors are controlled in part by circulating male-specific proteins produced by the fat body (FB) [20], a specialized tissue that performs energy storage functions equivalent to mammalian liver and white adipose tissue. Finally, sex differences in carbohydrate metabolism in cells of the adult intestine are controlled by signaling from the male gonad and couple diet with sperm production [21].

For any species, less is known about intrinsic, tissue-specific sex differences during development (reviewed in [4]). For those differences that are manifested before animals are sexually mature, it is not clear if they impact lifespan, healthspan and/or reproduction later in life. We previously identified and characterized how the SPEN family of RNA-binding proteins in *Drosophila* –– Split ends (Spen) and Spenito (Nito) –– act in the larval FB to maintain proper fat levels [22,23]. A connection between sex determination and fat storage came from parallel findings that Nito is also required for proper sex determination via the canonical *Sex-lethal* (*Sxl*) pathway [24-28]. As a first step to understanding intrinsic sex differences during *Drosophila* development, here we measure metabolic differences in larvae –– which are sexually immature –– and explore roles for Nito and the Sxl pathway in these differences.

## Results and Discussion

### *Metabolic sexual dimorphism in* Drosophila *larvae*

The larval developmental stage precedes pupae and adults. We first compared larval body fat using a density-based assay [23]. Animals of the commonly used *w*^*1118*^ experimental genetic background were sorted by sex and analyzed separately. Significant sexual dimorphism was observed, with males having higher overall fat levels than females (Fig 1A). Intriguingly, this sex difference is opposite to that found in adults [6,29,30]. The adult dimorphism develops gradually over time: fat levels are equivalent in newly-eclosed males and females [5]. Female adults consume more food than males, with mated females eating even more [10,18,31]. We found no significant sex difference in larval food consumption or locomotor activity (Fig 1B and C), suggesting that these behaviors do not contribute to the observed differences in fat levels. To ask if fat dimorphism at the organismal level reflects dimorphism at the molecular level, we performed lipidomic analysis on whole male or female larvae. Indeed, partial least squares-discriminant analysis (PLS-DA) revealed clustering in lipidomic profiles of male and female larvae (Fig S1). Males had higher levels of acylcarnitines, diacylglycerols, and several triacylglycerols (Fig 1D, Fig S1 and Table S1), consistent with our indirect body fat results from the density assay.

**Figure 1.**
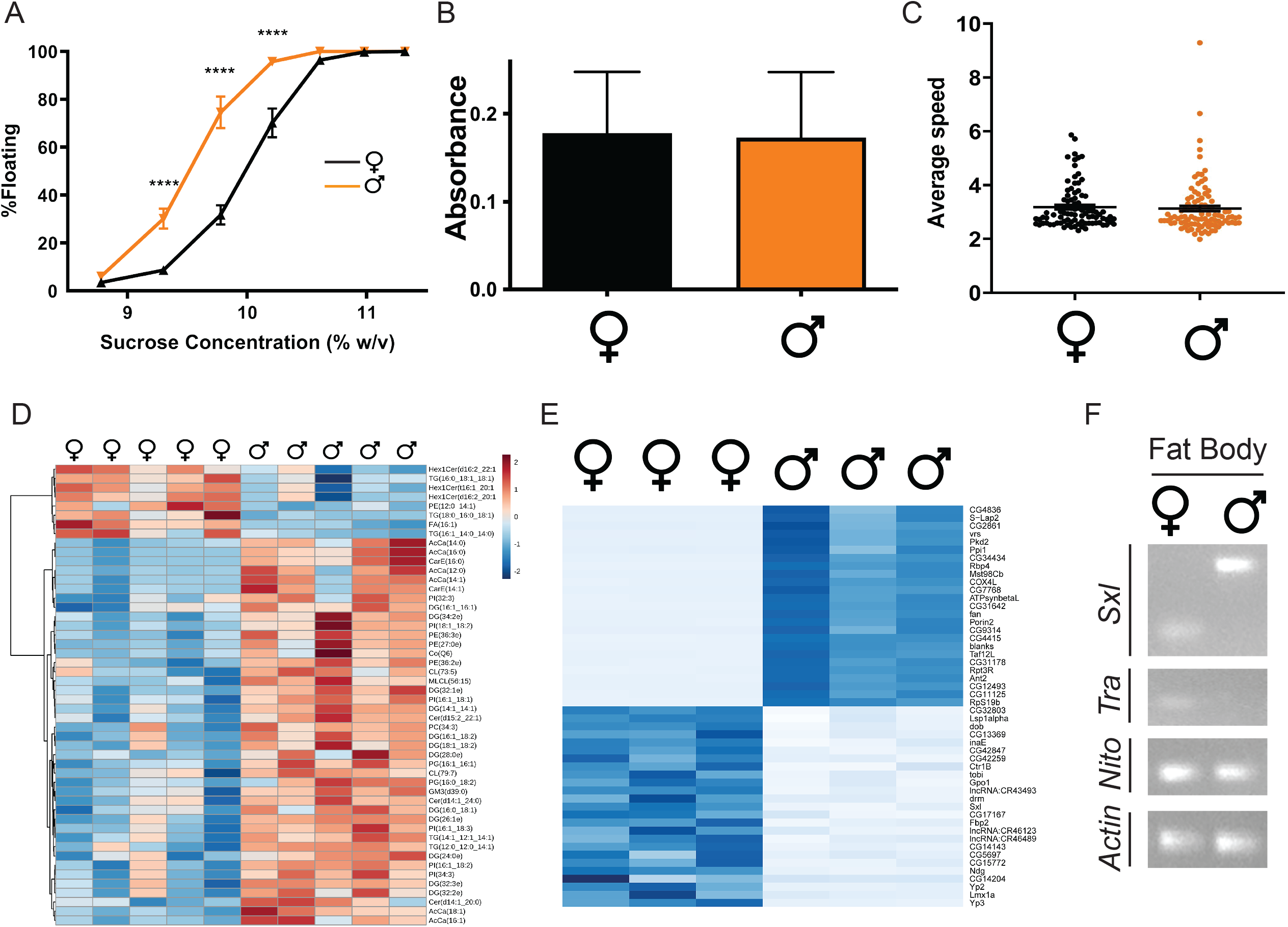
Sexual dimorphism in metabolism and metabolic gene expression. (**A**) Percent floating male (black) or female (orange) larvae in increasing sucrose densities (percent weight/volume). n=9 biological replicates per sample, 50 larvae per replicate. P values represent results from two-way ANOVA (****P < 0.0001). Error bars represent SEM. (**B**) Absorbance at 520 nm as a measure of food intake. n=4 biological replicates per sample, 20 larvae per replicate. Error bars represent SD. (**C**) Average larval speed, pixels/sec. n=4 biological replicates per sample, 15 larvae per replicate. Error bars represent SEM. (**D**) The heat map shows the top 50 most significant lipidomic differences between male and female larvae. Darker shades of orange indicate higher levels, darker shades of blue indicate lower levels of the specified lipid. n=5 biological replicates per sample,10 larvae per replicate. (**E**) Heatmap representing RNA-seq results of male versus female FB depicting the top 25 differentially upregulated and downregulated genes ranked by adjusted P value and ordered by log2 fold-change values. Darker shades of blue indicate higher expression. n=3 biological replicates. (**F**) RT-PCR products representing the indicated transcripts in RNA extracted from male or female larval fat bodies were separated on 2% agarose gels. *Actin* is a loading control. n=3 biological replicates; shown is a representative experiment.

In parallel, we isolated RNA from FBs from male and female larvae and performed RNA sequencing (RNA-seq). Among the 200 genes most significantly up-or down-regulated were multiple genes with known or predicted roles in lipid metabolism, including *Glycerophosphate oxidase 1* (*Gpo1*), the acylglycerolerol lipase *inactivation no afterpotential E* (*inaE*), and the lipase family genes *CG5162* [32], *doppelganger von brummer* (*dob*) [33], *Yolk protein 2* (*Yp2*), and *Yolk protein 3* (*Yp3*) [34] (Fig 1E, Fig S2, and Table S2). Yp3 was the most strongly downregulated transcript in male FBs (Fig 1E and Table S2). In the vitellogenesis process, yolk proteins synthesized in the FBs of adult females are secreted into circulation and ultimately taken up by developing oocytes to become the major protein components of yolk (reviewed in [35]). Following fertilization, the energy stored in yolk fuels embryogenesis. Yolk proteins lack conserved residues required for lipase activity [36] but are major components of the lipid droplet proteome [37] and have been proposed to transport lipids to oocytes [38]. Yp3 is among the most abundant larval proteins [39] and was the most strongly downregulated larval hemolymph protein upon starvation [40], further pointing to a role in larval organismal energy balance prior to oogenesis.

Expression of genes involved in glycolysis, oxidative phosphorylation and fatty acid metabolism was also dimorphic. Among the 200 genes most significantly up-or down-regulated were *Phospholipase A2 group III* (*GIIIspla2*), a predicted pyruvate kinase (*CG12229*), *Cytochrome c oxidase subunit 4-like* (*COX4L*), *ATP synthase, β subunit-like* (*ATPsynβL*), the long-chain-fatty-acid-CoA ligase *heimdall* (*hll*) [41], and the fatty acid elongase *Baldspot* [42] (Table S2). Also significant were *Carnitine palmitoyltransferase 2* (*CPT2*) [43] and *fatty acid binding protein* (*fabp*) (Table S2). Functional enrichment analysis using g:Profiler [44] revealed significant (P<0.05) sex-specific enrichment for multiple pathways relevant to the lipidomics changes we observed: Fatty acid beta-oxidation, Glycolysis/Gluconeogenesis, Carbon metabolism, and Pyruvate metabolism (Table S2). These differences in gene expression are fully consistent with the observed increase in body fat in males.

Notably, we saw little overlap with published adult sex-specific differences in metabolic genes [5] (Table S2), consistent with the phenotypic contrast with adults, where females are fatter. In our RNA-seq data we noticed sex-specific FB expression of *Sxl* and a downstream sex determination gene, *transformer* (*tra*) (Fig 1E). RT-PCR confirmed presence of the female-specific *Sxl* isoform exclusively in the female FB, and the male-specific isoform exclusively in the male FB (Fig 1F). We interpret these results as evidence that the canonical sex-determination pathway operates in larval FB cells. We note that in the larval stages, the gonads are embedded in the FB [45,46], but if sex-specific expression differences were restricted to the gonad, we would have seen mixed expression signals from FB cells. Male gonads in the larva are much bigger (>3-fold) than female gonads [45,46]. Thus we speculate that, analogous to the extra stores adult females require to support ovarian development [47], male larvae might need more stored energy for gonad growth and development.

### Nito regulates metabolic sexual dimorphism and sex determination in the fat body

Due to Nito’s role in sex determination, and because we observed sex-specific expression of sex-determinant genes in FB cells, we tested whether FB-specific depletion of Nito via RNAi alters the dimorphic expression of the sex determination gene *Sxl*. As expected, RNAi control animals expressed the respective male and female transcripts in the FB (Fig 2A). However, upon Nito depletion we observed the male *Sxl* isoform in female FBs (Fig 2A). These expression patterns are consistent with equivalent effects of Nito depletion in the wing [27] and with a masculinization of the female FB in the absence of Nito, and support the hypothesis that Nito is required in FB cells to establish and/or maintain a sex-specific larval FB identity.

**Figure 2.**
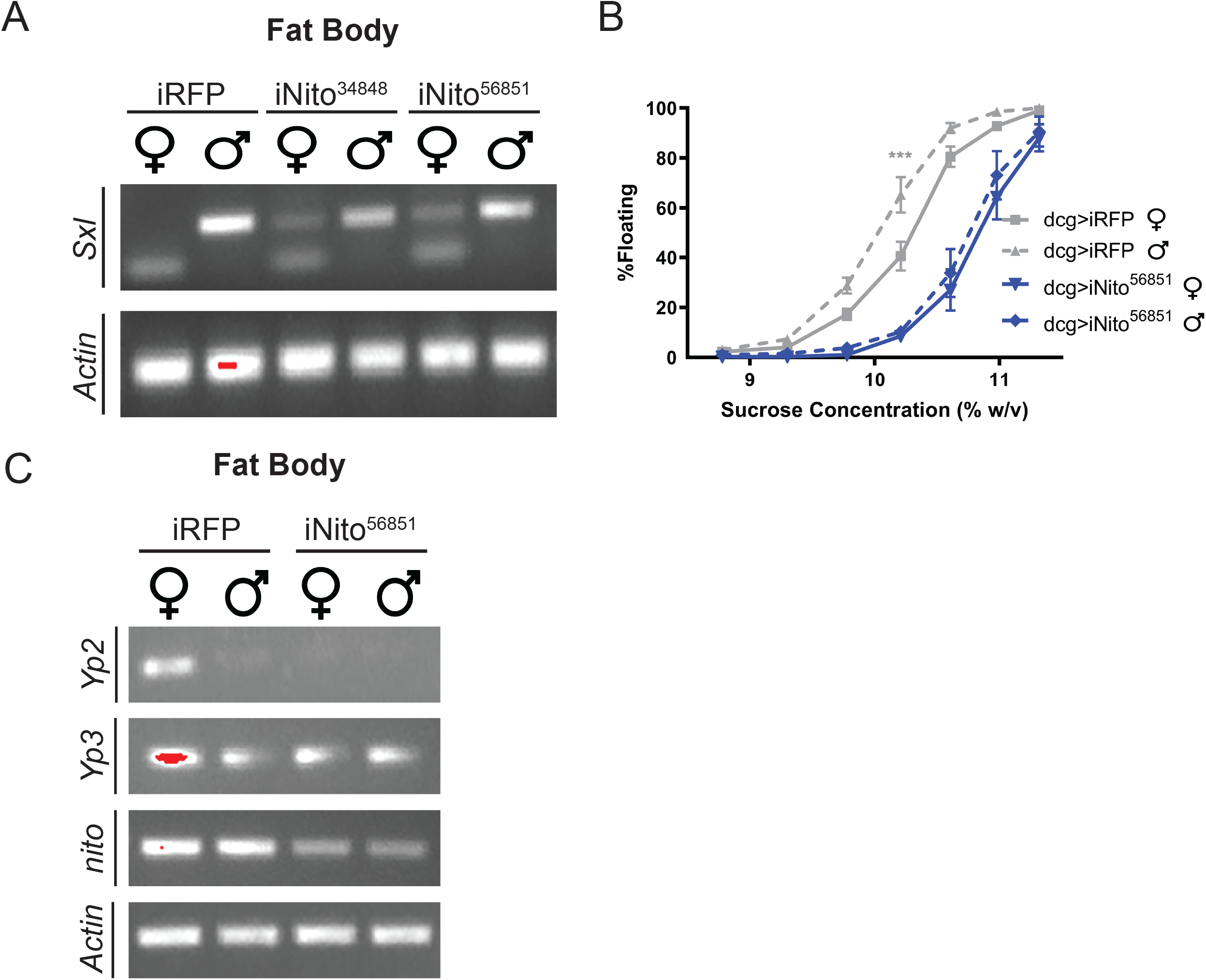
Nito is required for metabolic sexual dimorphism in the fat body. (**A**,**C**) RT-PCR products representing the indicated transcripts in RNA extracted from iNito and iRFP KD male and female larval fat bodies were separated on 2% agarose gels. *Actin* is a loading control. n=3 biological replicates; shown is a representative experiment. (**B**) Percent floating larvae in increasing sucrose densities (percent weight/volume). FB-specific Nito KD (dcg>iNito, blue) compared to KD control (dcg>iRFP, gray). Females are shown in solid lines and males in dashed lines. n=8 biological replicates per sample, 50 larvae per sample. P values represent results from two-way ANOVA, ***P < 0.001. Error bars represent SEM.

Using mixed-sex measurements, we previously found that FB depletion of Nito alters larval body fat [22]. We repeated this analysis but separated larvae by sex. Larvae with Nito-depleted FBs were lean, but there was no longer a significant difference between males and females (Fig 2B). At the molecular level, depleting Nito also eliminated the sex differences we had identified by RNA-seq in levels of transcripts encoding metabolic enzymes. Specifically, in confirmation of our RNA-seq data, RT-PCR revealed increased levels of *Yp2* and *Yp3* in control females compared to males (Fig 2C). By contrast, levels of *Yp2* and *Yp3* transcripts were similar in the Nito-depleted FB of males and females (Fig 2C). These results are consistent with a role for Nito in promoting metabolic sexual dimorphism in the FB via differential expression of genes controlling metabolism.

### Sex determination pathway effects on larval body fat

Our results indicate that metabolic sexual dimorphism is intrinsic to the FB. To ask directly if dimorphic gene expression in fat cells is sufficient to dictate sex-specific fat storage, we overexpressed the female determinant isoform of Tra (TraF) in male and female FBs. Consistent with Tra acting downstream of Sxl, there was no change in *Sxl* splicing or *nito* transcript levels (Fig 3A). Strikingly, expression of TraF in FB of males and females resulted in leaner males and females than the GFP overexpression control (Fig 3B). Consistent with metabolic feminization of males, following TraF overexpression we observed increased levels of *Yp2* and *Yp3*, similar to females (Fig 3C). Sex-specific *Yp2* and *Yp3* expression in adults is known to require *tra* function [48]. Our data show that this is also true in larval FBs. Tra was known to act in the FB to control dimorphism of larval body size via non-cell-autonomous insulin-like peptide signaling [29]. In adults, Tra promotes fat storage in females via control of hormone release from neurons [49]. We interpret our data as evidence of Tra-dependent dimorphism in metabolic gene expression intrinsic to the larval FB.

**Figure 3.**
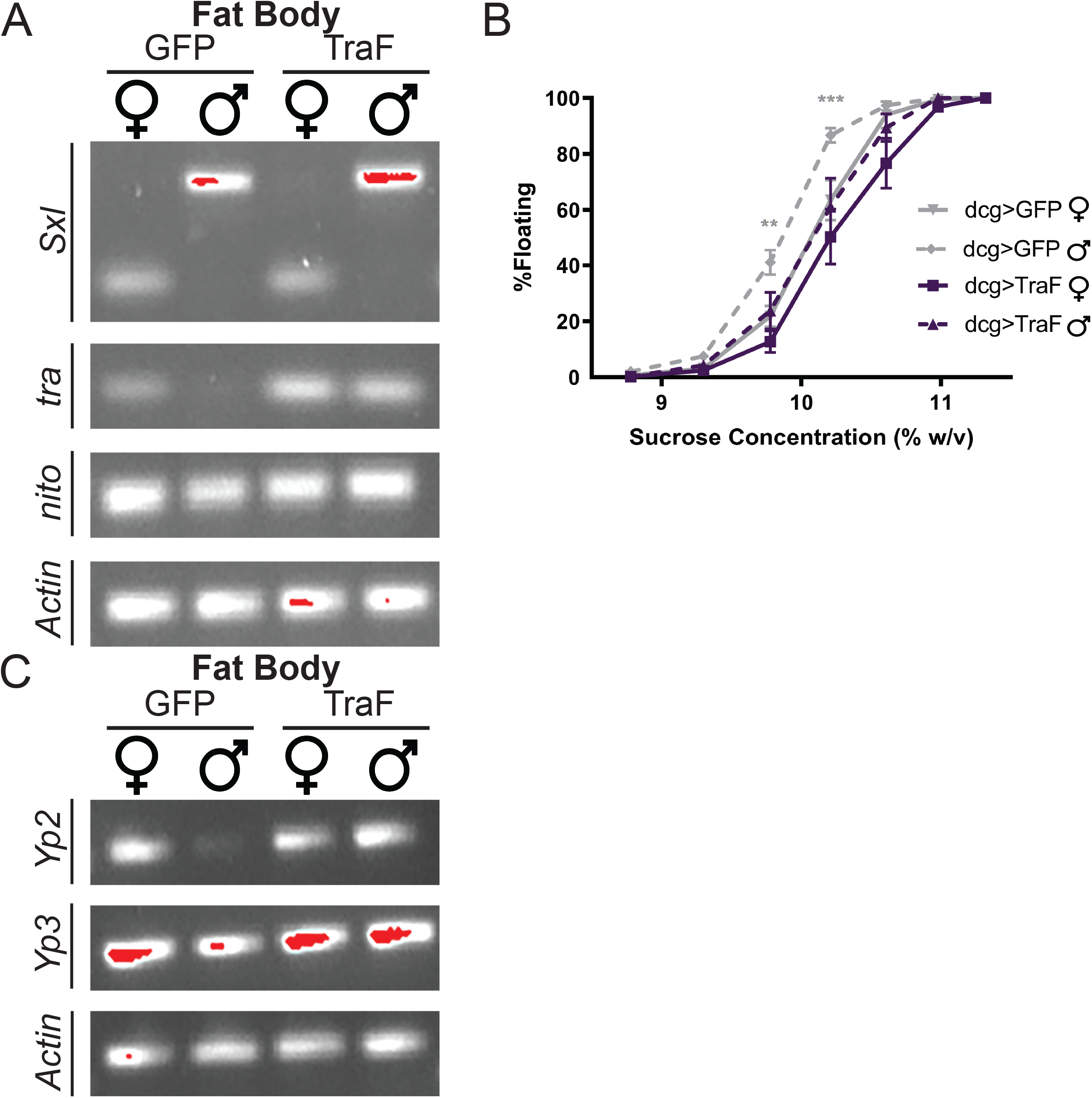
Fat Body-specific TraF expression feminizes metabolism in male larvae. (**A**,**C**) RT-PCR products representing the indicated transcripts in RNA extracted from TraF- and GFP-overexpressing male and female larval fat bodies. *Actin* is a loading control. n=3 biological replicates; shown is a representative experiment. (**B**) Percent floating larvae in increasing sucrose densities (percent weight/volume). FB-specific TraF overexpression (dcg>TraF, purple) compared to overexpression control (dcg>GFP, gray). Females are shown in solid lines and males in dashed lines. n=9 biological replicates per sample, 50 larvae per sample. P values represent results from two-way ANOVA, **P < 0.01, ***P < 0.001.. Error bars represent SEM.

Nito depletion and TraF overexpression both resulted in lean phenotypes and collapse of dimorphism. However, and as expected, FB-specific Nito depletion masculinized *Yp2* and *Yp3* expression in females and TraF overexpression feminized *Yp2* and *Yp3* expression in males (Fig 2C and 3C). The lean phenotype observed upon Nito depletion is stronger than that following TraF expression (Fig 2B and 3B). We previously characterized Nito’s antagonistic role to Spen function in regulating fat levels and showed that Spen is not required in the FB for metabolic dimorphism [22]. We therefore propose that Nito is required in two parallel pathways: one that regulates metabolism in a sex-specific manner and – – together with its sibling, Spen –– another that regulates metabolism in a sex-independent manner.

Given Nito’s role in RNA modification with N^6^-methyladenosine (m^6^A) as part of the canonical sex determination pathway [19-23], we predict that Nito-dependent m^6^A modification of transcripts encoding key metabolic enzymes results in dimorphic expression and ultimately metabolic differences. Indeed, m^6^A RNA modification in mice is an essential regulator of sex-specific differences in lipid metabolism [50]. High levels of m^6^A modification are present on lipogenic mRNAs in mice liver, with males having higher levels than females [50]. Additionally, loss of m^6^A in males leads to “feminization” of lipid composition [50]. It is not known which m^6^A targets control fat storage and how different components of the m^6^A machinery contribute to this regulation.

Taken together, our data raise new questions, demanding a deeper understanding of how overall organismal dimorphic differences result from a balance between intrinsic genetic versus hormonal differences and to what extent these differences influence healthspan and reproduction. Understanding these questions becomes even more important in the context of exogenous introduction of sex hormones and/or hormone blockers, such as hormonal therapies used as cancer treatments or as gender-affirming healthcare services.

## Materials and methods

### Fly Strains and husbandry

*w*^*1118*^ (Bloomington *Drosophila* Stock Center stock number (BL) 3605), *w*^*1118*^; *dcg > GAL4* (BL 7011), *y*^*1*^*sc v*^*1*^*sev*; *UAS-Nito*-RNAi (BL 56851), *y*^*1*^ *v*^*1*^; *UAS-Nito-*RNAi (BL 56860), *y*^*1*^*sc v*^*1*^*sev*; *UAS-RFP-*RNAi (BL 67852), *w*^*1118*^;*UAS-GFP* (BL 4775) and *w*^*1118*^; *UAS*-*TraF* (BL 4590) were obtained from the Bloomington *Drosophila* Stock Center. All lines were backcrossed to the *w*^*1118*^ stock. Unless otherwise specified, all animals were reared at 25°C, 60% humidity and fed a modified Bloomington media (1 L: yeast 15.9 g, soy flour 9.2 g, yellow cornmeal 67.1 g, light malt extract 42.4 g, agar 5.3 g, light corn syrup 90 g, propionic acid 4.4 ml, Tegosept (Apex Bioresearch Products #20-258, 380 g in 1L 100% ethanol) 8.4 mL. Experimental media (1 L: yeast 35g, soy flour 9.2 g, yellow cornmeal 65 g, light malt extract 42.4 g, agar 5.3 g, light corn syrup 70 mL, propionic acid 4.4 mL, Tegosept 8.4 ml). was made fresh each week and used for no longer than one week. Crosses were made with 100-120 virgin females with 50-60 males. Eggs were collected for 5 hours on grape plates at 25°C, 60% humidity and 50 first-instar larvae were transferred 22-24 hrs later into a vial with experimental media.

### Density Assay

For sexed density assays, 50 wandering-third-instar larvae of each sex were sorted per sample prior to the assay. Density assays were then performed as previously described [23,51]. For each experiment, genetic background controls were also tested by crossing either the siblings of each UAS-line or the driver line with *w*^*1118*^. The resulting male and female progeny showed similar dimorphism to controls and there were no significant effects of the insertions on density. All experimental conditions and genotypes were analyzed with 8-9 independent samples. ANOVA was used to calculate statistical significance with GraphPad Prism software.

### RT-PCR

Total RNA was extracted from 50 third-instar FBs dissected from sexed larvae using 500 µL of TRIzol (Ambion Cat # 15596018), and purified using the Direct-zol Miniprep Plus kit digested with DNase I (Zymo Research Cat # R2072). Total purified RNA was used for reverse transcription using SuperScript IV Reverse transcriptase (Invitrogen Cat # 18090010). Semi-quantitative PCR was performed using *Taq* DNA polymerase with standard *Taq* buffer (New England BioLabs Cat # M0273S). PCR products were analyzed on 2% Agarose gels with 0.5 ng/L Ethidium bromide using a 1kb Plus DNA Ladder (New England BioLabs Cat # N3200S) for size reference. Primers are listed below. Forward primer (_F), reverse primer (_R).

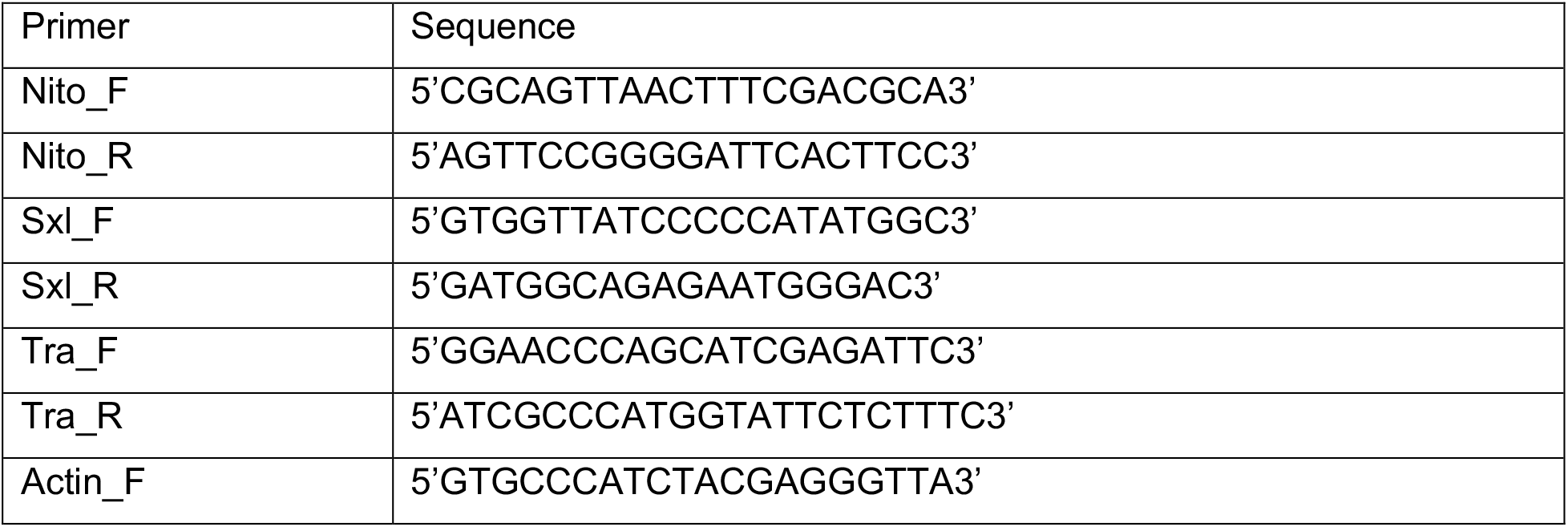
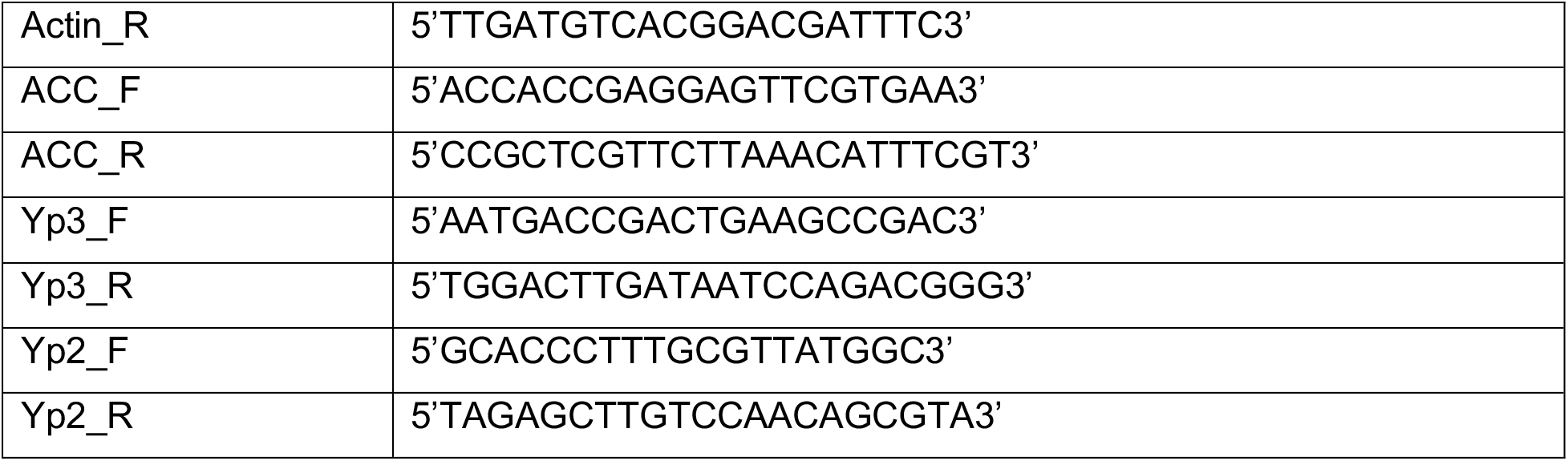

### Feeding Assay

Twenty sexed, early third-instar larvae were used per sample to measure intake of yeast containing 0.5% food dye (FD&C Red #40) on an agar plate at 25°C for 30 min, as previously described [23]. Four independent biological samples were analyzed by ANOVA using GraphPad Prism software.

### Activity Assay

Fifteen sexed, pre-wandering third-instar larvae were collected and tracked for movement as previously described [52]. Four independent samples were analyzed by Mann-Whitney and Kolmogorov-Smirnov tests using GraphPad Prism software.

### Lipidomics

#### Sample preparation

Lipids were extracted via a protein crash modified from a previously described method [52,53]. Wildtype whole larvae (n=10, biological experiment repeated 5 times per each condition (male vs female)), were homogenized and extracted at 15 mg/mL ratio in ice-cold methanol. Following homogenization, samples were vortexed 30 minutes, followed by centrifugation at 12700 RPM for 10 minutes at 4°C. 100 μL of supernatant was transferred to a new autosampler tube for sample analysis.

#### UHPLC-MS data acquisition and processing

Analyses were performed as previously published [54]. Briefly, the analytical platform employs a Vanquish UHPLC system (Thermo Fisher Scientific, San Jose, CA, USA) coupled online to a Q Exactive mass spectrometer (Thermo Fisher Scientific, San Jose, CA, USA). Lipid extracts were resolved over an ACQUITY HSS T3 column (2.1 × 150 mm, 1.8 µm particle size (Waters, MA, USA) using an aqueous phase (A) of 25% acetonitrile and 5 mM ammonium acetate and a mobile phase (B) of 90% isopropanol, 10% acetonitrile and 5 mM ammonium acetate. For negative mode analysis the chromatographic the gradient was as follows: 0.3 mL/min flowrate and 30% B at 0 minutes, 0.3 mL/min flowrate and 100% B at 3 minutes, 0.3 mL/min flowrate and 100% B at 4.2 minutes, 0.4 mL/min flowrate and 30% B at 4.3 minutes, 0.4 mL/min flowrate and 30% B at 4.5 minutes, 0.3 mL/min flowrate and 30% B at 5 minutes. For positive mode analysis the chromatographic gradient was as follows: 0.3 mL/min flowrate and 10% B at 0 minutes, 0.3 mL/min flowrate and 95% B at 3 minutes, 0.3 mL/min flowrate and 95% B at 4.2 minutes, 0.45 mL/min flowrate and 10% B at 4.3 minutes, 0.4 mL/min flowrate and 10% B at 4.5 minutes, 0.3 mL/min flowrate and 10% B at 5 minutes. The Q Exactive mass spectrometer (Thermo Fisher) was operated in positive ion mode, scanning in Full MS mode (2 μscans) from 150 to 1500 m/z at 70,000 resolution, with 4 kV spray voltage, 45 sheath gas, 15 auxiliary gas. When required, dd-MS2 was performed at 17,500 resolution, AGC target = 1e5, maximum IT = 50 ms, and stepped NCE of 25, 35 for positive mode, and 20, 24, and 28 for negative mode. Calibration was performed prior to analysis using the Pierce™ Positive and Negative Ion Calibration Solutions (Thermo Fisher Scientific).

#### Data analysis

Acquired data was converted from raw to mzXML file format using Mass Matrix (Cleveland, OH, USA). Samples were analyzed in randomized order with a technical mixture injected interspersed throughout the run to qualify instrument performance. Lipidomics data were analyzed using LipidSearch 4.0 (Thermo Scientific), which provides lipid identification on the basis of accurate intact mass, isotopic pattern, and fragmentation pattern to determine lipid class and acyl chain composition. Graphs, heat maps and statistical analyses (T-Test) and Partial Least Squares-Discriminant Analysis (PLS-DA) were performed using MetaboAnalyst 5.0 [54].

### RNA sequencing

#### Sample preparation

Total RNA was extracted from 50 third-instar FBs dissected from sexed larvae using 500 µL of TRIzol (Ambion Cat # 15596018), and purified using the Direct-zol Miniprep Plus kit digested with DNase I (Zymo Cat # R2072). RNA sequencing and library prep was performed at the University of Colorado Anschutz medical campus Genomics Core. Libraries were prepped according to the manufacturer’s protocol using the Universal Plus mRNA-Seq library preparation kit with NuQant (TECAN Cat # 0520-24).

#### Data acquisition and processing

Libraries were sequenced with an Illumina NovaSEQ 6000 system. Reads were filtered and trimmed using Trim Galore! [55] (version 0.6.5). Reads were aligned to the fly genome using HISAT [56] (version 2.2.1). Mapped reads were sorted using SAMtools [57] and expression was quantified using featureCounts [58] (version 1.11). DESeq2 [59] (version 1.34.0) was used for differential expression analysis. Pathway analysis was performed using gProfiler [44] (version 0.2.1). All code relating to this project will be available at https://github.com/rnabioco/reis-fly-fat-body upon publication.

## Data availability statement

Fly lines are available upon request. Supplemental files available at FigShare. File S1 shows principal component analysis. File S2 shows heatmaps of gene expression changes. Table S1 has lipidomics data analysis. Table S2 has gene expression data analysis. Raw gene expression data are available at GEO with the accession number: XXXX.

## Figure legends

**Fig S1.**
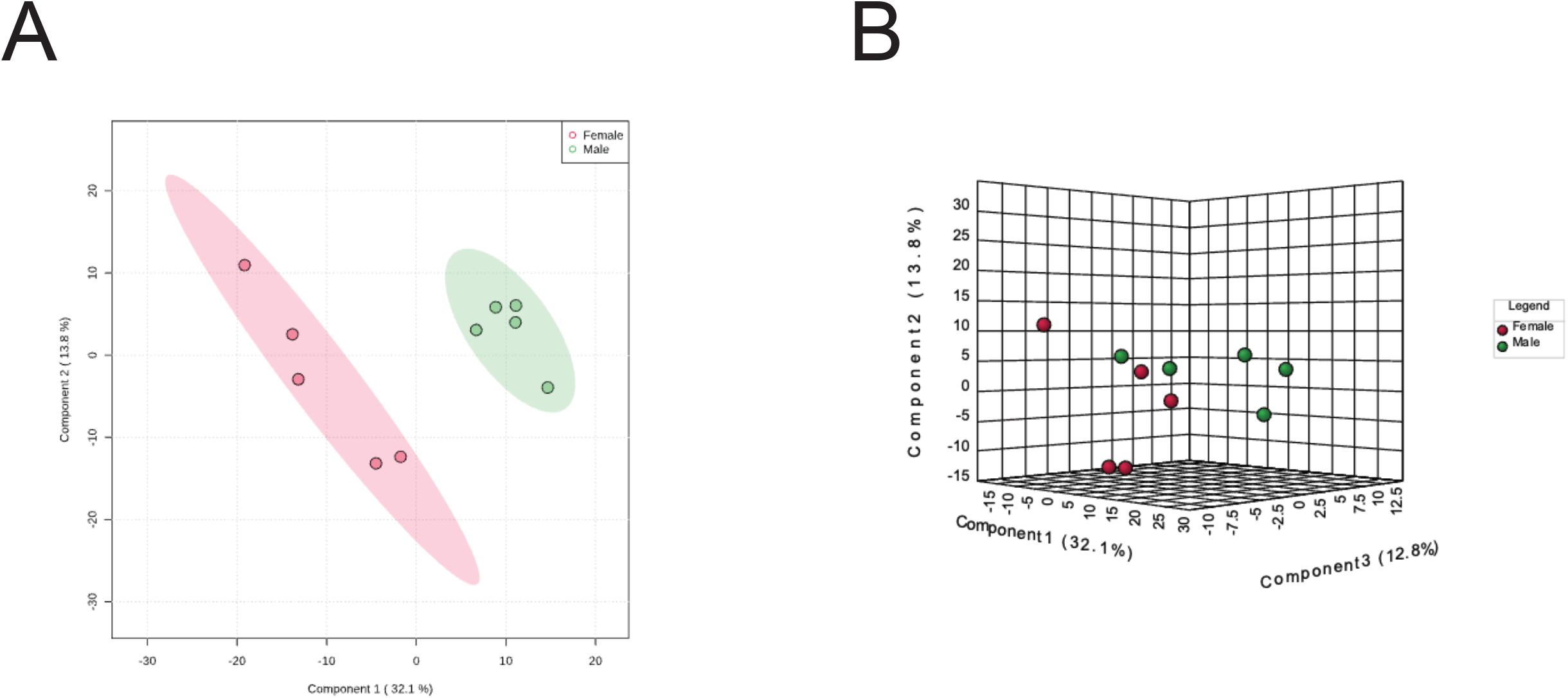
Partial least squares-discriminant analysis (PLS-DA) reveals clustering in lipidomic profiles of male and female larvae. (A) The lipidomic differences of male versus female larvae as a 2D PLS-DA plot (B) The lipidomic differences of male versus female larvae as a 3D PLS-DA plot.

**Table S1.Lipidomic analysis of male and female larvae.**

**Fig S2.**
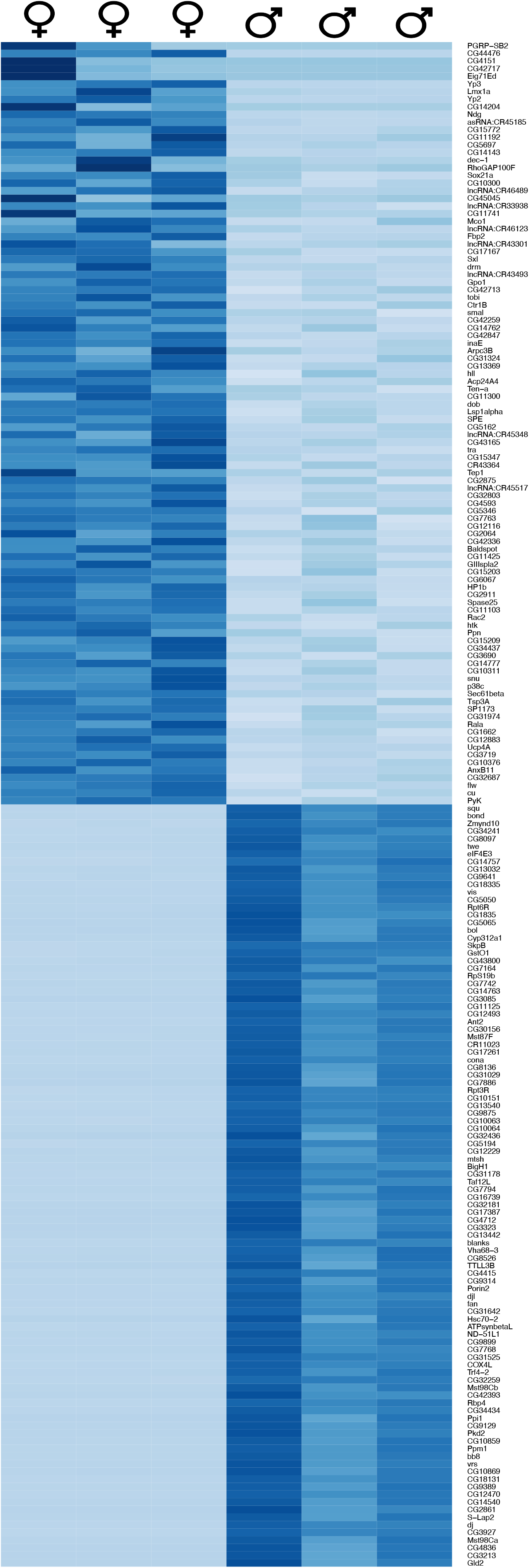
Heatmap of top differentially expressed genes in male versus female fat bodies. Heatmap depicting the top 100 differentially upregulated and downregulated genes by adjusted P value and ordered by log2 fold-change values. Darker shades of blue indicate higher expression.

**Table S2.RNA-seq analysis of male and female larvae.**

## Acknowledgements

We would like to thank the Bloomington Stock Center for generously maintaining and providing fly stocks and Michael McMurray for his helpful comments and editing. We also thank Laura George, who performed preliminary studies on the dimorphism of Oregon R larvae. A.V.D. is supported by NIH-T32-GM136444, the Victor W. and Earleen D. Bolie Graduate Scholarship, and a CU Anschutz RNA Bioscience Initiative Scholar award. T.R. is supported by the National Institute of Diabetes and Digestive and Kidney Diseases (R01DK106177) and a Pilot Award from the RNA Biosciences Initiative of the University of Colorado Anschutz Medical Campus.

